# APOBEC-mediated mutagenesis in urothelial carcinoma is associated with improved survival, mutations in DNA damage response genes, and immune response

**DOI:** 10.1101/123802

**Authors:** Alexander P. Glaser, Damiano Fantini, Kalen J. Rimar, Joshua J. Meeks

**Affiliations:** Northwestern University, Department of Urology, Chicago, IL, 60607

**Author notes:** **Corresponding author**: Joshua J. Meeks, MD PhD, 303 E. Chicago Ave., Tarry 16-703, Chicago, IL 60611. **Abbreviations and Acronyms** TCGA: The Cancer Genome Atlas ssDNA: single stranded DNA APOBEC: apolipoprotein B mRNA editing catalytic polypeptide-like GCAC: Genome Data Analysis Center MAF: mutation annotation format “APOBEC-high”: tumors enriched for APOBEC mutagenesis “APOBEC-low”: tumors not enriched for APOBEC mutagenesis.

**Keywords:** Urinary bladder neoplasms, APOBEC Deaminases, Mutagenesis, DNA damage, Interferon

## Abstract

**Background:** The APOBEC family of enzymes is responsible for a mutation signature characterized by a TCW>T/G mutation. APOBEC-mediated mutagenesis is implicated in a wide variety of tumors, including bladder cancer. In this study, we explore the APOBEC mutational signature in bladder cancer and the relationship with specific mutations, molecular subtype, gene expression, and survival. We hypothesized that tumors with high levels of APOBEC-mediated mutagenesis would be enriched for mutations in DNA damage response genes and associated with higher expression of genes related to activation of the immune system.

**Methods:** Gene expression (n=408) and mutational (n=395) data from the Cancer Genome Atlas (TCGA) bladder urothelial carcinoma provisional dataset was utilized for analysis. Tumors were split into “APOBEC-high” and “APOBEC-low” tumors based on APOBEC enrichment score. Analysis was performed with R.

**Findings:** Patients with APOBEC-high tumors have better overall survival compared to those with APOBEC-low tumors (38.2 vs 18.5 months, p=0.005). Tumors enriched for APOBEC mutagenesis are more likely to have mutations in DNA damage response genes (*TP53, ATR, BRCA2*), and chromatin regulatory genes (*MLL, MLL3*), while APOBEC-low tumors are more likely to have mutations in *FGFR3* and *KRAS. APOBEC3A* and *APOBEC3B* expression correlates with total mutational burden, regardless of bladder tumor molecular subtype. APOBEC mutagenesis and enrichment is associated with increased expression of immune-related genes, including interferon signaling.

**Interpretation:** Tumors enriched for APOBEC mutagenesis are more likely to have mutations in DNA damage response genes and chromatin regulatory genes, potentially providing more single-strand DNA substrate for *APOBEC3A* and *APOBEC3B*, leading to a hypermutational phenotype and the subsequent immune response.

**Highlights:**

- ABPOEC enzymes, particularly *APOBEC3A* and *APOBEC3B*, are responsible for the predominant pattern of mutagenesis in bladder cancer
- Tumors enriched for APOBEC-mediated mutagenesis are more likely to have mutations in DNA damage response genes and chromatin regulatory genes, while tumors not enriched for APOBEC-mediated mutagenesis are more likely to have mutations in *KRAS* and *FGFR3*
- APOBEC enrichment is associated with upregulation of genes involved in the immune response

## 1. Introduction

Urothelial carcinoma has one of the highest mutation rates of any sequenced cancer to date ^1^. High-throughput next generation sequencing analyses such as The Cancer Genome Atlas (TCGA) and others have identified a mutational signature characterized by a TCW>T/C mutation thought to be attributable to the apolipoprotein B mRNA editing catalytic polypeptide-like (APOBEC) family of enzymes ^1-3^. This mutational pattern is the predominant pattern in in bladder cancer and is also frequently found in breast, cervical, head and neck, and lung cancers ^1,3,4^.

The APOBEC family consists of 11 members, including *AID, APOBEC1, APOBEC2, APOBEC3A, APOBEC3B, APOBEC3C, APOBEC3D, APOBEC3F, APOBEC3G, APOBEC3H*, and *APOBEC4*. These enzymes function as cytosine deaminases and are involved in C>U deamination in single-stranded DNA (ssDNA), and likely function physiologically in antiretroviral defense ^5-8^. However, in tumor cells, these enzymes are likely responsible for hypermtuation at cytosine bases in exposed ssDNA, known as kataegis ^9^. The APOBEC3 family, and particularly *APOBEC3A* and *APOBEC3B* ^5,10-13^ but also *APOBEC3H*^*14*^, are the predominant APOBEC enzymes theorized to contribute to cancer mutagenesis.

Several studies have linked *APOBEC3B* expression with mutagenesis ^13,15^, but expression alone does not fully explain this mutational signature, and *APOBEC3A* may also play a significant role ^10^. In breast cancer, DNA replication stress and mutations in DNA repair genes have been linked to APOBEC-mediated mutagenesis ^16^, potentially due to increased availability of ssDNA substrate for enzymatic deamination ^17,18^. However, less is known about the downstream effects of APOBEC mutagenesis in bladder cancer.

In this study, we explore the APOBEC mutational signature in bladder cancer and its relationship with specific mutations, molecular subtype, gene expression, and survival. We hypothesized that tumors with high levels of APOBEC-mediated mutagenesis would be enriched for mutations in DNA damage response genes and express genes related to activation of the immune system at higher levels.

## 2. Methods

### 2.1 The Cancer Genome Atlas data

The Cancer Genome Atlas (TCGA) bladder urothelial carcinoma data was downloaded from the Broad Institute Genome Data Analysis Center (GDAC) (http://gdac.broadinstitute.org)^19^ and from cBioPortal (http://cbioportal.org)^20^. Data from GDAC was downloaded on November 8, 2016, from the analysis timestamp “analyses_2016_01_28” (doi:10.7908/C19G5M58)^21^. Downloaded data includes clinical and demographic data (age, sex, tumor stage, overall survival), mutation annotation files (MutSig 2CV v3.1; MAF file; Mutsig_maf_modified.maf.txt) and mRNA expression (Immumina HiSeq RNAseqV2). RNA-seq mRNA expression levels are presented as RNA-seq by expectation-maximization (RSEM) values ^22^.

Clinical information was available on 412 samples, RNA-seq data was available on 408 samples, and mutation information was available on 395 samples for TCGA BLCA data version 2016_01_28. Overlap between the 412 patients with clinical information, 408 patients with RNA-seq data, and 395 patients with mutation annotation information yields 391 patients. Three outliers were removed from mutational analysis (TCGA-DK-A6AW, >150 mutations/Mb; TCGA-XF-AAN8 and TCGA-FD-A43, both with ≤5 total mutations).

### 2.2 Mutation analysis and APOBEC enrichment

Analysis and visualization of mutations from mutation annotation format (MAF) files was performed using R version 3.3.3, Bioconductor^23^ version 3.4 (http://www.Bioconductor.org), and MAFtools version 1.0.55 ^24^. Mutation rates per sample were calculated using MutSig2CV v3.1 from the Broad Institute GDAC ^1^. APOBEC enrichment score based on the frequency of TCW>T/G mutations was calculated as previously described ^3,4,25^. Samples were classified into two groups: “APOBEC-high” based on APOBEC enrichment > 2 and Benjamini-Hochberg^26^ false-discovery-rate corrected p-value < 0.05; and “APOBEC-low” based on an APOBEC enrichment < 2 and/or Benjamini-Hochberg false-discovery-rate corrected p-value ≥ 0.05. Survival outcomes between patients with APOBEC-high-enrichment and APOBEC-low-enrichment was performed using log-rank test and Kaplan-Meyer curves (R packages survival ν 2.41-2, survminer v0.3.1, ggplot2 v2.2.1).

Significantly differentially mutated genes between APOBEC-high-enrichment and APOBEC-low-enrichment groups was performed using MAFtools ^24^ as previously described ^27^ and visualized with oncoplots and forest plots.

### 2.3 Molecular Subtyping

Molecular subtyping of 408 bladder urothelial carcinoma samples with RNA-seq data was performed using R, Bioconductor, and multiClust version 1.4.0 ^28^. Samples were classified as luminal, p-53-like, basal, or claudin-low as previously described ^29^ with hierarchical clustering using Euclidean distance and Ward’s linkage method (ward.D2; hierarchical clustering analysis shown in **Supplementary Figure 1**). Differences in mutational load and expression of APOBEC3 enzymes (RSEM) between tumor subtypes was compared using ANOVA.

### 2.4 Differential gene expression and gene expression associated with APOBEC enrichment

Differential expression analysis between APOBEC-high and APOBEC-low tumors was performed using R, Bioconductor, limma version 3.30.13 and edgeR version 3.16.5 ^30,31^. Association of gene expression with numeric APOBEC enrichment score was performed using Spearman’s rho. Functional annotation of genes was performed with DAVID version 6.8 (http://david.ncifcrf.gov) ^32,33^ and visualized with Enrichr (http://amp.pharm.mssm.edu/Enrichr/) ^34,35^.

## 3. Results

### 3.1. APOBEC mutagenesis in bladder cancer

In bladder cancer, as the mutation load increases, the frequency of specific nucleotide conversions changes, and C>G mutations become more common (**Figure 1A**). Many of these mutations are a specific contextual TCW>T/G mutation attributed to the APOBEC family of enzymes. Of 389 tumors in the provisional TCGA bladder urothelial carcinoma dataset, 324 are enriched for APOBEC mutagenesis (“APOBEC-high”) vs. 64 with low or no enrichment (“ABPOEC-low”). APOBEC-high tumors have improved overall survival compared to APOBEC-low tumors (median overall survival 38.2 vs 18.5 months, p=0.0050, **Figure 1B**).

**Figure 1.**
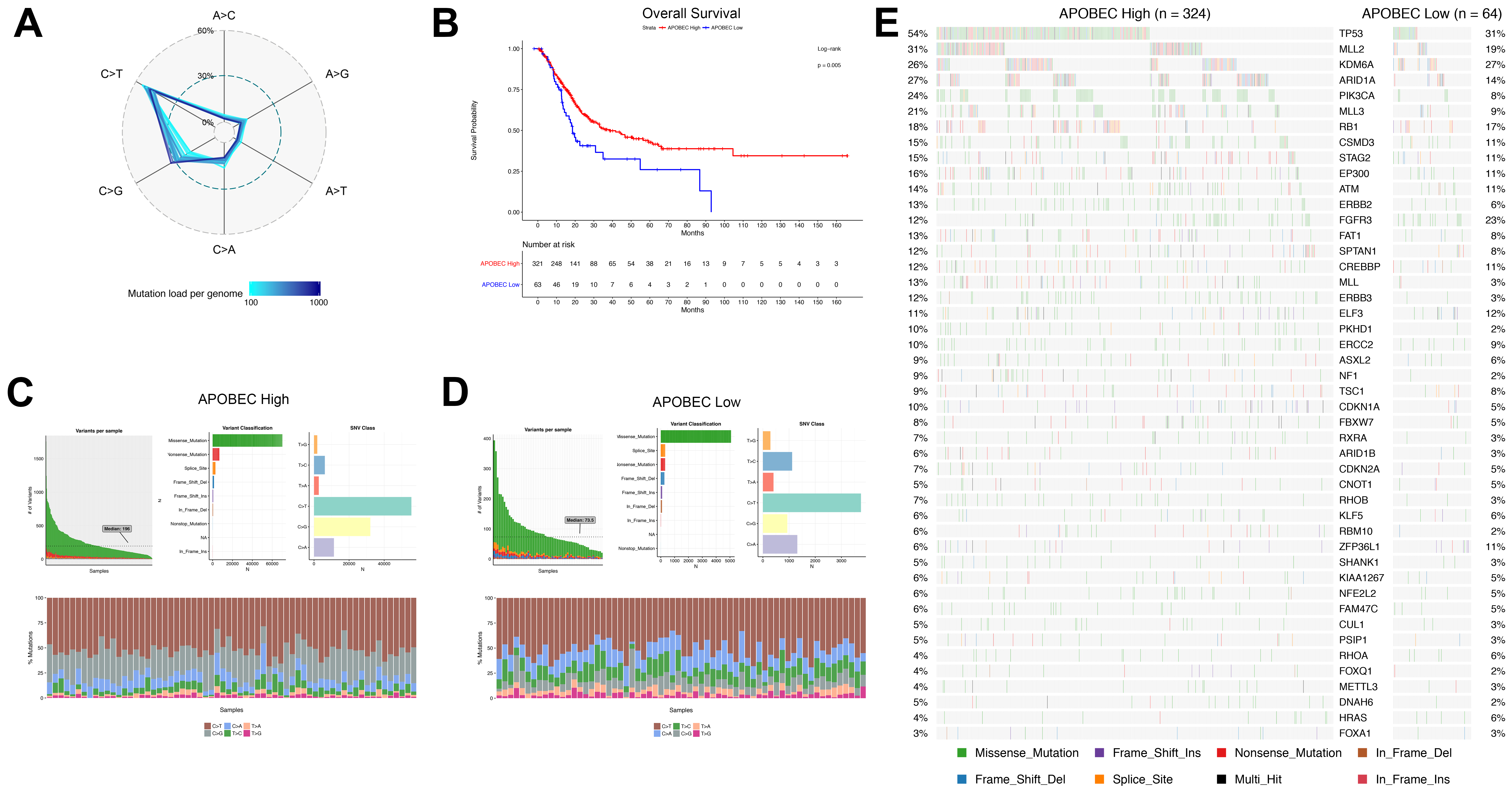
APOBEC-mediated mutagenesis in the TCGA bladder cancer cohort (n=389). (A) Percentage of single nucleotide variations (SNVs) as a function of mutation load. Genomes were binned in groups of 20 samples according to mutation load. (B) Kaplan-Meier survival curve of APOBEC-high and APOBEC-low bladder tumors. (C) Summary of mutagenesis in APOBEC-high tumors, including number of variants per sample, variant classification, and class of SNV. Stacked barplot demonstrates percentage of SNVs in a representative sample of 64 tumors for comparison with APOBEC-low tumors. (D) Summary of mutagenesis in APOBEC-low tumors, including number of variants per sample, variant classification, class of SNV, and stacked barplot of the percentage of SNV per tumor. (E) Oncoplot of the top genes commonly mutated in bladder cancer in APOBEC-high and APOBEC-low tumors.

APOBEC-low tumors are more likely to be low-grade (17% vs 3%, p<0.0001), but there is no significant difference in age at diagnosis, tumor stage, tumor subtype, or patient smoking history category between groups (**Supplementary Table 1**). Interestingly, a higher frequency of Asian patients was noted in the APOBEC-low group (26% vs 7%), and *APOBEC3B* was expressed at a lower level in Asian patients vs. patients not of Asian ethnicity (**p<0.0001, Supplementary Figure 2**).

APOBEC-high tumors have a higher number of variants per sample and a higher proportion of C>T and C>G mutations (**Figure 1C-D**). Despite an association of APOBEC enrichment score with total mutations, 42% of APOBEC-high tumors have a mutational burden below the median (**Supplementary Figure 3**), and APOBEC-high and APOBEC-low tumors share many of the commonly mutated genes in bladder cancer (**Figure 1E**).

### 3.2. Frequency of mutations in APOBEC-high and APOBEC-low tumors

To determine what mutations were associated with APOBEC mutagenesis, we next compared significantly mutated genes between APOBEC-high and APOBEC-low tumors. After correction for multiple comparisons, APOBEC-high tumors were more likely to have mutations in *TP53, PIK3CA, ATR, BRCA2, MLL, MLL3*, and *ARID1A*, among others; while APOBEC-low tumors were more likely to have mutations in *KRAS* and *FGFR3* (Figure 2A-B; complete list of differentially mutated genes in **Supplementary Table 2**).

**Figure 2.**
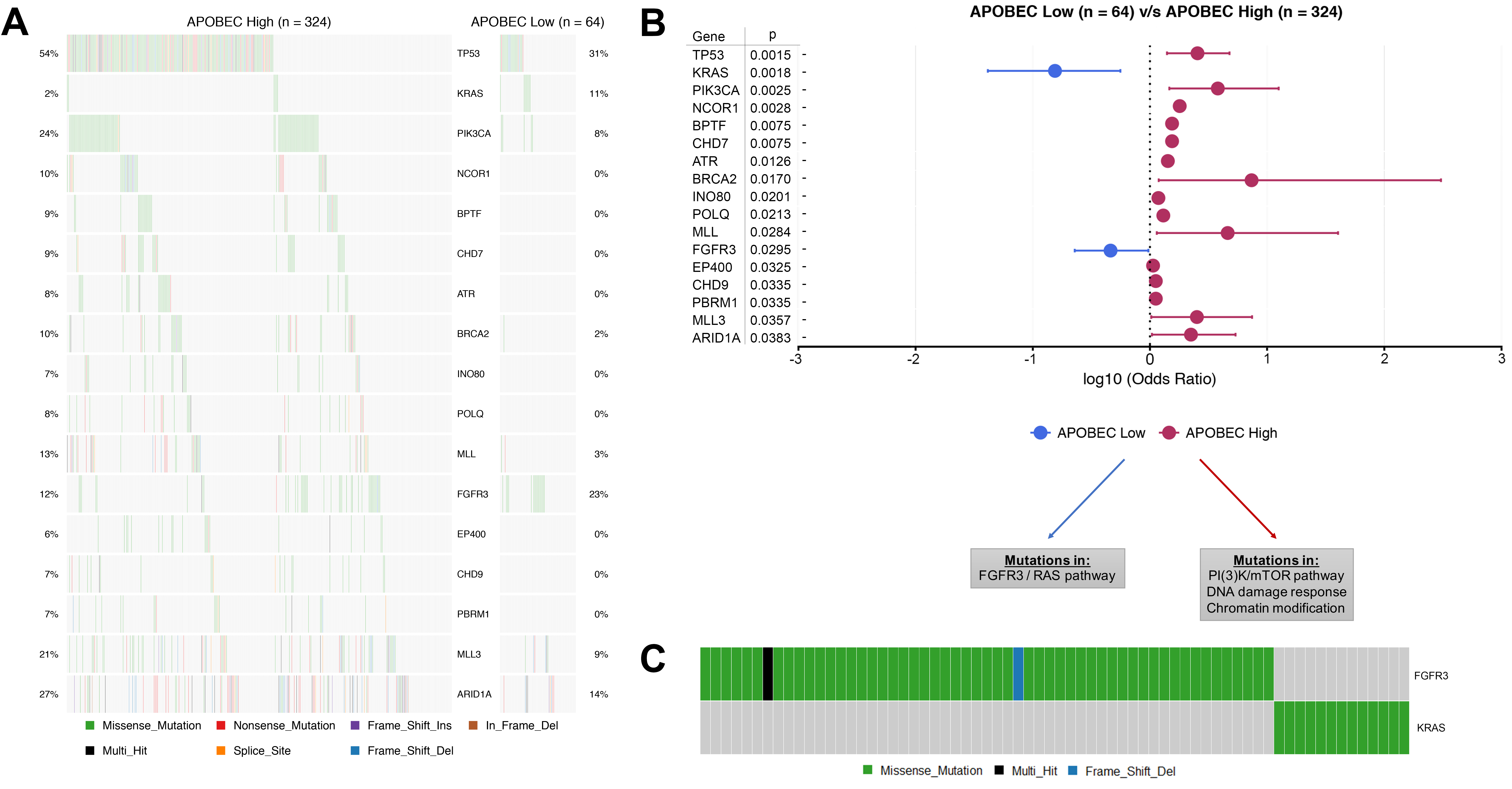
Significantly differentially mutated genes in APOBEC-high and APOBEC-low tumors. (A) Oncoplot of genes differentially mutated in APOBEC-high and APOBEC-low tumors. (B) Forestplot of differentially mutated genes in APOBEC-high and APOBEC-low tumors with log10 odds ratio and 95% confidence intervals and adjusted p-value (MutSig 2CV 3.1; MAFtools v1.0.55). (C) Oncoprint of nonsynonymous mutations in *FGFR3* and *KRAS* (n=395).

Functional annotation of differentially mutated genes demonstrates that APOBEC-high tumors are enriched for mutations in DNA damage repair genes (e.g. *TP53, ATR, BRCA2, POLQ*) and chromatin modification genes including *MLL, MLL3, AIRD1A, NCOR1, BPTF, CHD7*, and others. A full list of gene ontology terms associated with genes mutated in APOBEC-high tumors can be found in **Supplementary Table 3**. APOBEC-low tumors were significantly more likely to harbor mutations in *KRAS* and *FGFR3*. As in the original TCGA dataset (n=131) ^36^, non-synonymous mutations in *KRAS* and *FGFR3* are mutually exclusive (**Figure 2C**).

### 3.3. Expression of APOBEC3A and APOBEC3B correlate with mutational burden

We next investigated the role of expression of APOBEC3 enzymes with total mutations in both the entire TCGA bladder cancer dataset and in the four molecular subtypes (luminal, p53-like, basal, and claudin-low). Expression of *APOBEC3A* and *APOBEC3B* correlate with the total mutation burden in bladder cancer (**Figure 3A-B**), while expression of *APOBEC3F* and *APOBEC3G* correlate weakly with total mutations and *APOBEC3C, APOBEC3D*, and *APOBEC3H* do not correlate with total mutations (**Supplementary Figure 4**).

**Figure 3.**
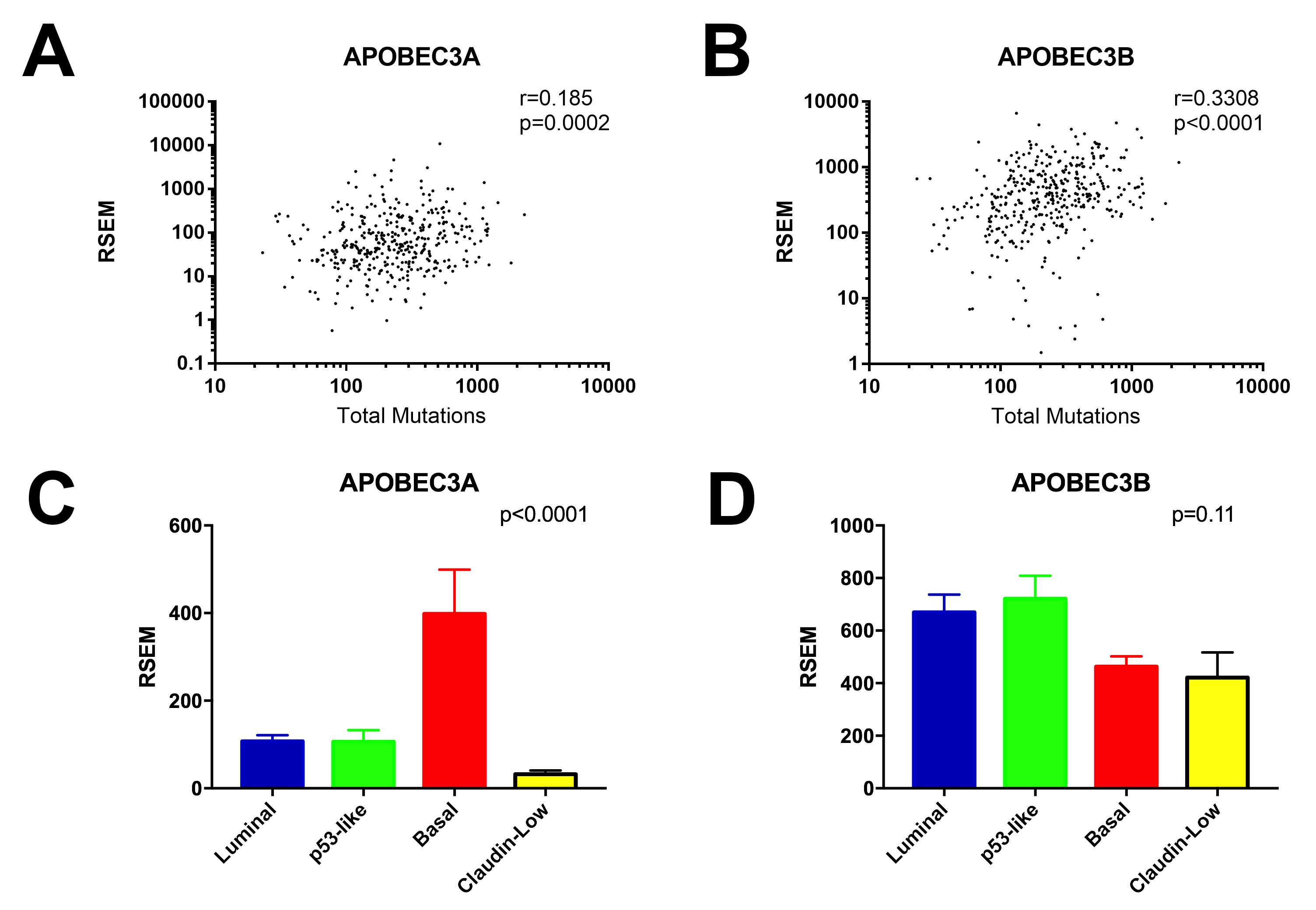
Correlations of *APOBEC3A* and *APOBEC3B* expression with mutational burden and molecular subtype. (A) Spearman correlation between total mutations and *APOBEC3A* expression. (B) Spearman correlation between total mutations and *APOBEC3B* expression. (C) Expression of *APOBEC3A* in the molecular subtypes of bladder cancer. (D) Expression of *APOBEC3B* in the molecular subtypes of bladder cancer.

Total mutations, APOBEC enrichment score, and the percentage of tumors classified as APOBEC-high vs APOBEC-low are not different between subtypes of bladder cancer (**Supplementary Figure 5**). However, *APOBEC3A* is expressed at a significantly higher level in the basal subtype than in luminal, p53-like, or claudin-low subtypes (**Figure 3C**), while *APOBEC3B* is evenly expressed across subtypes (**Figure 3D**). Despite this, *APOBEC3A* expression levels correlate with total mutations in every subtype, as do *APOBEC3B* expression levels (**Supplementary Figure 6**).

### 3.4. Gene expression associated with APOBEC enrichment

We approached gene expression association with APOBEC enrichment by two methods. We first analyzed differentially expressed genes between APOBEC-high and APOBEC-low tumors. APOBEC-high tumors were enriched for expression of genes related to regulation of the immune response and lymphocyte-mediated immunity (**Figure 4A** **and Supplementary Table 4**), whereas APOBEC-low tumors with higher expression of genes related to transcription and translation (**Figure 4B** **and Supplementary Table 5**). Similarly, correlation between numeric APOBEC enrichment score and gene expression revealed a positive relationship between APOBEC enrichment score and gene families involved in IFN-gamma signaling, antigen presentation, and regulation of the immune response, including immune checkpoint HAVCR2 (also known as TIM-3; r=0.229, p<0.0001) (**Figure 4C** **and Supplementary Table 6**), while genes inversely correlated with APOBEC enrichment score were enriched in gene families related to transcription and translation (**Figure 4D** **and Supplementary Table 7**).

**Figure 4.**
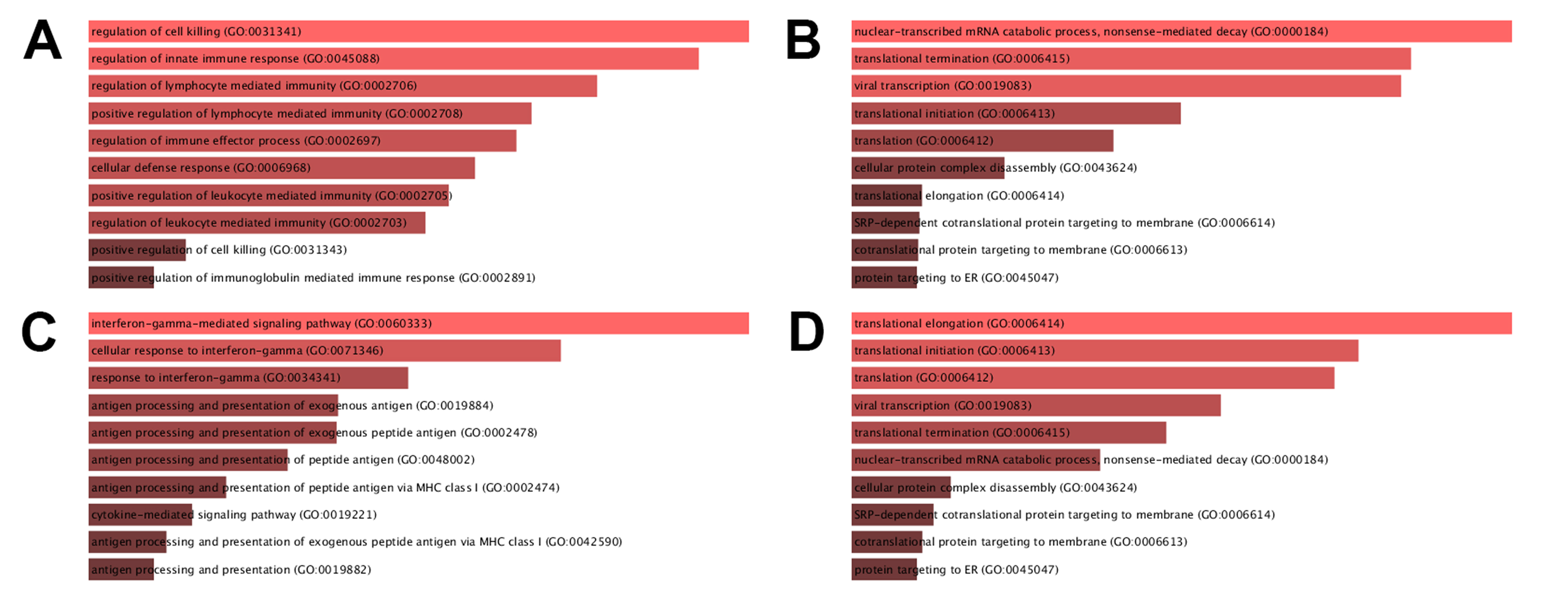
Barplot of gene ontology biological processes for: (A) genes highly expressed in APOBEC-high tumors (B) genes highly expressed in APOBEC-low tumors (C) genes positively correlated with APOBEC enrichment score (D) genes negatively correlated with APOBEC enrichment score. Figure generated with Enrichr.^34,35^ Bar size based on combined score based on p-value and deviation from expected rank.

## 4. Discussion

APOBEC mutagenesis is the predominant mutational pattern in bladder cancer. In this paper, we demonstrate that tumors enriched for APOBEC mutagenesis (APOBEC-high tumors) have better survival and are more likely to have mutations in DNA damage repair genes and chromatin regulation genes. Expression of *APOBEC3A* and *APOBEC3B* correlates with overall mutation load in bladder cancer, regardless of molecular subtype. In addition, APOBEC enrichment is associated with upregulation of immune-related genes including interferon signaling. Unexpectedly, we also demonstrate that tumors not featuring the APOBEC mutational pattern (APOBEC-low tumors) are significantly more likely to harbor mutations in *FGFR3* and *KRAS*, which are mutually exclusive.

Our work is consistent with several prior studies linking APOBEC expression to mutational burden. *APOBEC3B* expression is upregulated in breast cancer and is associated with overall mutations ^15^. Our group also demonstrated the relationship between APOBEC3B expression and mutations in bladder cancer^37^. Overexpression of *APOBEC3B* in breast cancer cell lines results in DNA fragmentation, increased C>T mutations, delayed cell cycle arrest, and eventual cell death ^15^. Furthermore, knockdown of *APOBEC3B* with short hairpin RNA in breast cancer cell lines decreases total number of uracil lesions, *TP53* mutations, and C>T mutations ^15^.

*APOBEC3A* expression was not initially detectable in breast cancer cell lines ^15^, and *APOBEC3B* expression correlates strongly with overall mutations in multiple malignancies ^13^,^15^, leading many to believe that APOBEC3B is responsible for the majority of these mutations. However, *APOBEC3A* expression is also correlated with mutational burden ^3^, as we demonstrate again here, *APOBEC3A* is highly proficient at cytidine hypermutation and creation of DNA double-strand breaks ^38,39^, and may have a larger role in mutagenesis than previously recognized ^10^.

Expression alone of APOBEC enzymes is insufficient to explain APOBEC-mediated mutagenesis. In this study, we demonstrate that although the basal subtype of bladder cancer has significantly higher expression of *APOBEC3A* than other subtypes, basal tumors do not have greater number of mutations or APOBEC enrichment compared to the other subtypes, and *APOBEC3A* expression correlates with total mutation burden even in subtypes with lower levels of *APOBEC3A* expression. This suggests that a baseline level of APOBEC expression is required for APOBEC mutagenesis, but above a certain threshold other factors are also influential ^4^. Post-translational modification regulating APOBEC enzymatic activity may play a role ^40^. Alternatively, as with any enzymatic reaction, increasing substrate availability may move equilibrium toward the accumulation of mutated DNA products.

The proposed substrate for APOBEC mutagenesis is ssDNA, a common DNA repair intermediate that may accumulate in cells with defects in DNA repair pathways ^41^. Interestingly, APOBEC-high tumors are more likely to have mutations in genes related to DNA repair and chromatin regulation. In breast cancer, tumor cell lines with high levels of *APOBEC3B* expression are more likely to have mutations in *TP53* ^*15*^. Furthermore, APOBEC-high breast cancers are more likely to have mutations in *TP53, NCOR1, MLL3* and other genes involved in DNA replication stress ^25^. In this study, we demonstrate that APOBEC-high bladder tumors are more likely to have mutations in *TP53, NCOR1, MLL3 (KMT2C), MLL (KMT2A), ATR, BRCA2*, and other genes related to DNA repair and chromatin modification.

We also demonstrated a higher frequency of *PIK3CA* mutations in APOBEC-high bladder tumors. *PIK3CA* has been previously reported to be mutated at a high frequency in specific TCW-containing helical motifs across a number of tumor types ^42^. Our analysis supports these results, with the majority of *PIK3CA* mutations in APOBEC-high tumors occurring in the helical domain at E542 and E545; no mutations in the helical domain were seen in APOBEC-low tumors (**Supplementary Figure 7**). These specific mutations in E542 and E545 have also been reported as hotspot mutations in breast cancer ^43^.

Interestingly, APOBEC-low tumors in this study were more likely to have mutations in the oncogenes *FGFR3* and *KRAS*, which are mutually exclusive. This suggests that tumors not enriched for the APOBEC mutational pattern may be driven by oncogenes which may dysregulate cellular homeostasis via mechanisms that do not result in accumulation of ssDNA intermediates used as substrate for APOBEC mutagenesis.

APOBEC enrichment was associated with overall survival and expression of immune related genes. *APOBEC3A* expression measured by Nanostring has previously been associated PD-L1 expression on tumor-infiltrating mononuclear cells in bladder cancer ^44^. APOBEC mutagenesis is associated with overall mutational burden in bladder cancer, which likely reflects downstream neoantigen burden and subsequent immune response ^45-47^.

Based on our results and the above discussion, we propose a working model of mutagenesis and the immune response in bladder cancer (Figure 5), in which a urothelial cell acquires one or more driver mutation(s). Accumulation of mutations in *TP53, ATR, BRCA2*, and/or other DNA damage response genes or chromatin regulation genes may result in the accumulation of ssDNA substrate for to *APOBEC3A* and *APOBEC3B*, leading to a high level of APOBEC-mediated mutagenesis and a hypermutation phenotype. This hypermutation in turn leads to a large neoantigen burden and the subsequent immune response generated from this increase in neoantigens. In addition, APOBEC enzymes, especially *APOBEC3A*, may be induced and overexpressed in response to interferon ^4,8,39,48^, potentially causing a positive feedback loop. In contrast, other tumors with mutations in the *FGFR3/RAS* pathway or other oncogenes may not expose sufficient substrate ssDNA to APOBEC enzymes to undergo significant APOBEC mutagenesis. These tumors have poor survival, despite an enrichment for *FGFR3* mutations and low-grade tumors, which were classically considered more benign phenotypes.

**Figure 5.**
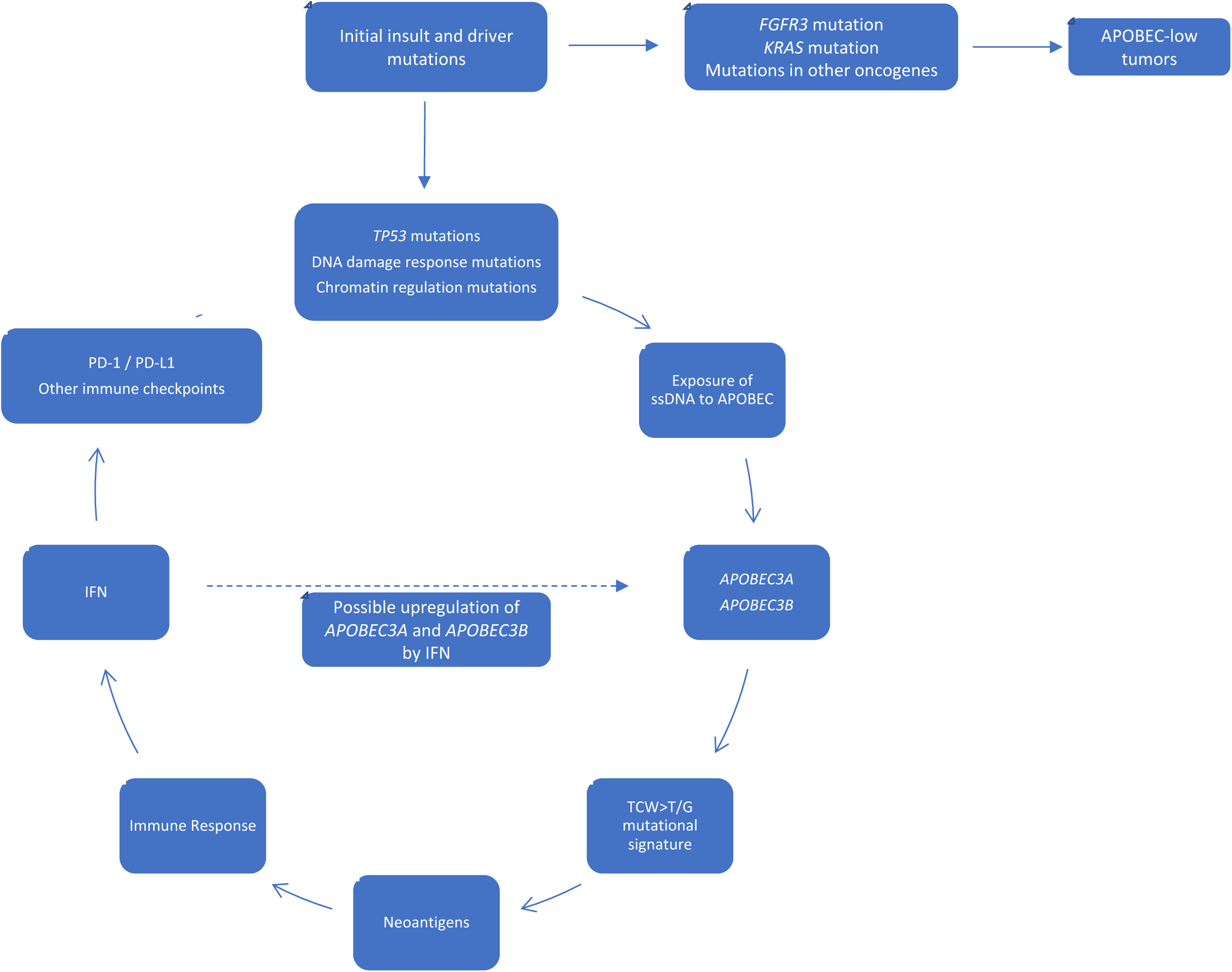
Working model of APOBEC-mediated mutagenesis in bladder cancer. Accumulation of mutations in *TP53, ATR, BRCA2*, and/or other DNA damage response genes or chromatin regulation genes may expose more substrate ssDNA to *APOBEC3A* and *APOBEC3B*, leading to a high level of APOBEC-mediated mutagenesis and a hypermutation phenotype, with subsequent neoantigen burden, immune response, and survival benefit. Tumors with mutations in *FGFR3* and *KRAS* may not expose enough substrate to APOBEC enzymes to promote APOBEC-mediated mutagenesis.

Several limitations of this study warrant mention. We utilized the provisional TCGA dataset, for which mutational and expression data is readily available. Results warrant replication in other datasets. The TCGA does not currently include any systemic treatment-related information. However, mutations in *ERCC2* ^*49,50*^ and other DNA repair genes ^51,52^ are associated with response to platinum-based therapy, and further investigation into the role of APOBEC mutagenesis and response to both cytotoxic chemotherapy and immunotherapy is warranted. Another limitation is the lack of a specific gene expression signature observed in APOBEC-low tumors other than transcription- and translation-related genes, potentially due to the heterogeneity of this group. In addition, gene expression correlations with APOBEC enrichment score in APOBEC-low tumors would not be expected to generate a strong signal, as these tumors by definition have a low and heterogeneous numerical APOBEC enrichment score.

In summary, APOBEC enzymes are a major source of mutation in bladder cancer. Tumors enriched for APOBEC mutagenesis have better survival and are more likely to have mutations in DNA damage repair genes and chromatin modifying genes. The APOBEC mutagenesis signature is associated with increased expression of immune-related genes. Bladder tumors not enriched for APOBEC mutagenesis are more likely to have mutations in *KRAS* and *FGFR3*, which are mutually exclusive, and these tumors have poor overall survival. Further study of the regulation of APOBEC enzymes, mutagenesis, and response to subsequent therapy may provide further insight into the mutational landscape and potential therapeutics for bladder cancer.

## 5. Acknowledgements

The results published here are in whole or part based upon data generated by the TCGA Research Network: http://cancergenome.nih.gov/. These results were presented in part at the American Urological Association 2017 meeting (Boston, MA; Abstract #17-3454).

## 8. Author Contributions

APG and JJM conceived the project ideas. JJM was responsible for project final design, supervision, interpretation of the data. APG, DF, and KR performed data collection and analysis. APG drafted the manuscript. All authors were involved in critical revision of the article and final approval of the version to be published.

